# MIMOSA2: A metabolic network-based tool for inferring mechanism-supported relationships in microbiome-metabolome data

**DOI:** 10.1101/2021.09.14.459910

**Authors:** Cecilia Noecker, Alexander Eng, Elhanan Borenstein

## Abstract

**Motivation:** Recent technological developments have facilitated an expansion of microbiome-metabolome studies, in which a set of microbiome samples are assayed using both genomic and metabolomic technologies to characterize the composition of microbial taxa and the concentrations of various metabolites. A common goal of many of these studies is to identify microbial features (species or genes) that contribute to differences in metabolite levels across samples. Previous work indicated that integrating these datasets with reference knowledge on microbial metabolic capacities may enable more precise and confident inference of such microbe-metabolite links.

**Results:** We present MIMOSA2, an R package and web application for model-based integrative analysis of microbiome-metabolome datasets. MIMOSA2 uses reference databases to construct a community metabolic model based on microbiome data and uses this model to predict differences in metabolite levels across samples. These predictions are compared with metabolomics data to identify putative microbiome-governed metabolites and specific taxonomic contributors to metabolite variation. MIMOSA2 supports various input data types and can be customized to incorporate user-defined metabolic pathways. We demonstrate MIMOSA2’s ability to identify ground truth microbial mechanisms in simulation datasets, and compare its results with experimentally inferred mechanisms in a dataset describing honeybee gut microbiota. Overall, MIMOSA2 combines reference databases, a validated statistical framework, and a user-friendly interface to facilitate modeling and evaluating relationships between members of the microbiota and their metabolic products.

**Availability and Implementation:** MIMOSA2 is implemented in R under the GNU General Public License v3.0 and is freely available as a web server and R package from www.borensteinlab.com/software_MIMOSA2.html.

## Background

Microbial community metabolism contributes to global nutrient cycling (1), metabolic dysregulation in human disease (2), and detoxification of pollutants (3), among other crucial processes. A growing number of studies investigate these processes by profiling the composition and metabolism of host-associated and environmental microbial communities using genomic and metabolomic technologies (4). Such studies commonly seek to answer two important questions: whether metabolic differences between environments (e.g. between healthy and diseased settings) can be attributed to differences in microbial composition and ecology, and if so, which specific microbial community members might be the key players generating such differences.

Broad surveys of microbial taxa and metabolites, which we refer to here as “microbiome-metabolome studies”, have great potential utility to answer these two questions by uncovering associations between taxa and metabolite abundances across samples. However, we have recently shown that calculating univariate correlations between taxa and metabolites may not identify mechanistic links between them with very high accuracy, depending on properties of the communities under study (5). Comparisons of taxon-metabolite associations with metabolite production in monoculture have also suggested a high false positive rate for this approach (6).

Some recently introduced tools address this challenge by using machine learning to infer complex models predicting metabolite abundances from microbial taxa (7–9). However, these can be difficult to interpret in terms of possible mechanisms, and in general do not account for or incorporate prior knowledge of microbial metabolism.

An alternative or complementary approach is to use metabolic reference databases such as the Kyoto Encyclopedia of Genes and Genomes (KEGG) (10), as well as collections of genome-scale metabolic reconstructions such as AGORA (11) and *embl_gems* (12), to generate and evaluate more precise hypotheses on the relationships between microbes and metabolites. We previously demonstrated that this conceptual approach can lead to new insights and released a preliminary but widely used R package (13), MIMOSA (Model-based Integration of Metabolite Observations and Species Abundances), for performing these analyses (14–19). Other methods to integrate microbiome and metabolome data with metabolic reference databases have also recently been introduced (20–23). To date, however, these tools all have some limitations in usability and/or compatibility with common data types and reference databases. Furthermore, none have been systematically evaluated for their ability to accurately infer direct mechanistic links between microbes and metabolites.

Here, we introduce MIMOSA2, an R package and web application (http://borensteinlab.com/software_MIMOSA2.html) for model-based integration of microbiome and metabolome data. MIMOSA2 uses metabolic reference databases to analyze paired microbiome and metabolite profiles, identifying metabolite differences that can be explained by specific microbiome features. MIMOSA2 expands and improves on the original proof-of-concept version of MIMOSA in several ways: First, it applies a new statistical algorithm for quantifying links between taxa and metabolites, which we have validated using simulated microbiome-metabolome data and experimental comparisons. It also implements several distinct strategies for constructing community metabolic models from different types of microbiome data and reference databases. Finally, it can be run either locally or via a web server with greatly improved ease of use, flexibility, and documentation.

## Implementation

MIMOSA2 integrates paired microbiome-metabolome datasets with reference reaction databases to generate specific mechanistic hypotheses. It constructs community metabolic models using microbiome compositional data and a reaction database, assesses whether measured metabolite concentrations are consistent with estimated community metabolic potential (CMP) across a set of samples, and identifies specific taxa and reactions that can explain observed metabolite variation (Figure 1). Below, we describe the analysis workflow steps and how to run an analysis using the web application.

**Figure 1.**
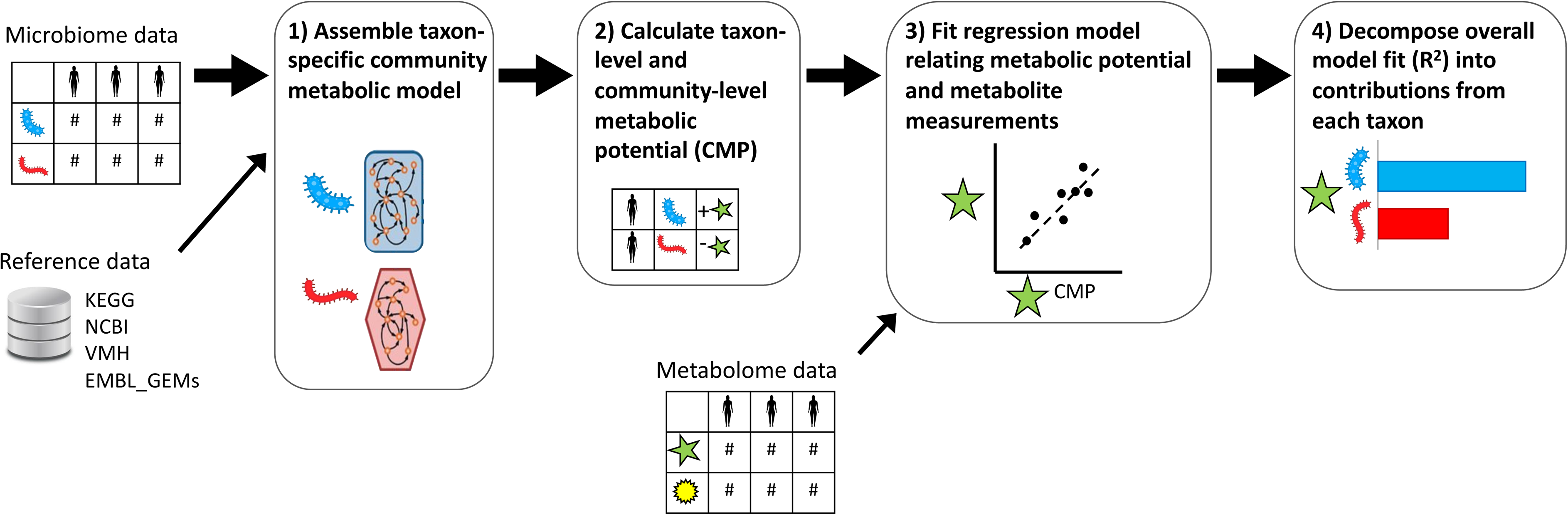
Summary of the MIMOSA2 analysis pipeline. In a MIMOSA2 analysis, microbiome data features are first linked to pre-processed reference databases to construct a community metabolic model describing the predicted metabolic reaction capabilities of each community member taxon (step 1). Next, this network model is combined with microbiome feature abundances to calculate community metabolic potential (CMP) scores for each metabolite, taxon, and sample, representing the approximate relative capacity to synthesize or utilize that metabolite (step 2). Total community metabolic potential scores for each metabolite are linked to metabolomics measurements using a regression model (step 3). For metabolites with a significant relationship between concentration and potential, the specific taxonomic contributors to each metabolite are then analyzed (step 4).

### Data input options

The basic input requirement for MIMOSA2 is a pair of datasets from the same set of samples: one of microbiome measurements and the other of metabolites. Each of these datasets may take a variety of forms, depending on the study’s design, experimental assays, and processing techniques.

Users can provide microbiome data generated from 16S rRNA or shotgun metagenomic sequencing studies. 16S rRNA data can be provided as a feature table of amplicon sequence variants (ASVs), or of closed-reference operational taxonomic units (OTUs) using either the GreenGenes or SILVA databases (24, 25). Data from a shotgun metagenomic study can be provided in the form of a table of KEGG Ortholog (10) functional abundances. Metagenomic data can be either unstratified (total KEGG Ortholog abundances in each sample) or stratified by microbial taxa (in the formats produced by either HUMAnN or PICRUSt2 software (26, 27)).

Metabolite data can be produced from any metabolomics platform, but it must contain putative metabolite identifications in the form of either metabolite names or KEGG compound IDs. If metabolite names are provided, they are mapped to KEGG IDs for the main analysis using the metabolite ID cross-referencing utility from the *MetabolAnalystR* package (28).

### Reference data and metabolic model construction

MIMOSA2 uses reference data to estimate how community metabolic potential varies across a set of microbiome samples. It takes advantage of genome-scale metabolic model data from multiple alternative sources to use the most appropriate reference data for a given dataset. Specifically, MIMOSA2 can currently generate a metabolic model based on one of three sources. First, it can use a curated set of reactions from the KEGG database. If OTU data is provided, KEGG reactions are inferred for each OTU using PICRUSt 1 (29) pre-computed outputs.

Alternatively, MIMOSA2 can link 16S rRNA amplicon data to one of two large collections of genome-scale metabolic reconstructions: the AGORA collection of genome-scale metabolic models of gut microbial species (v1.0.2) (11), or the *embl_gems* library of genome-scale metabolic models for all 5,587 reference and representative bacterial genomes in RefSeq (12).

The method used to map microbiome taxa abundances to metabolic reactions depends on the input data type (described fully in Methods and illustrated in Figure 6). 16S rRNA sequence variants are mapped using *vsearch* to either RNA genes for the AGORA genome collection (see Methods), or to Greengenes 99% OTU representative sequences. Greengenes OTUs are mapped to KEGG using the precomputed genome inferences from PICRUSt 1.1.3 (29), and are linked to AGORA models using a pre-calculated alignment between the two databases. KEGG Ortholog abundances provided from a shotgun metagenomic dataset or another method (such as from PICRUSt2 (30)) can be directly linked to KEGG reactions.

**Figure 6.**
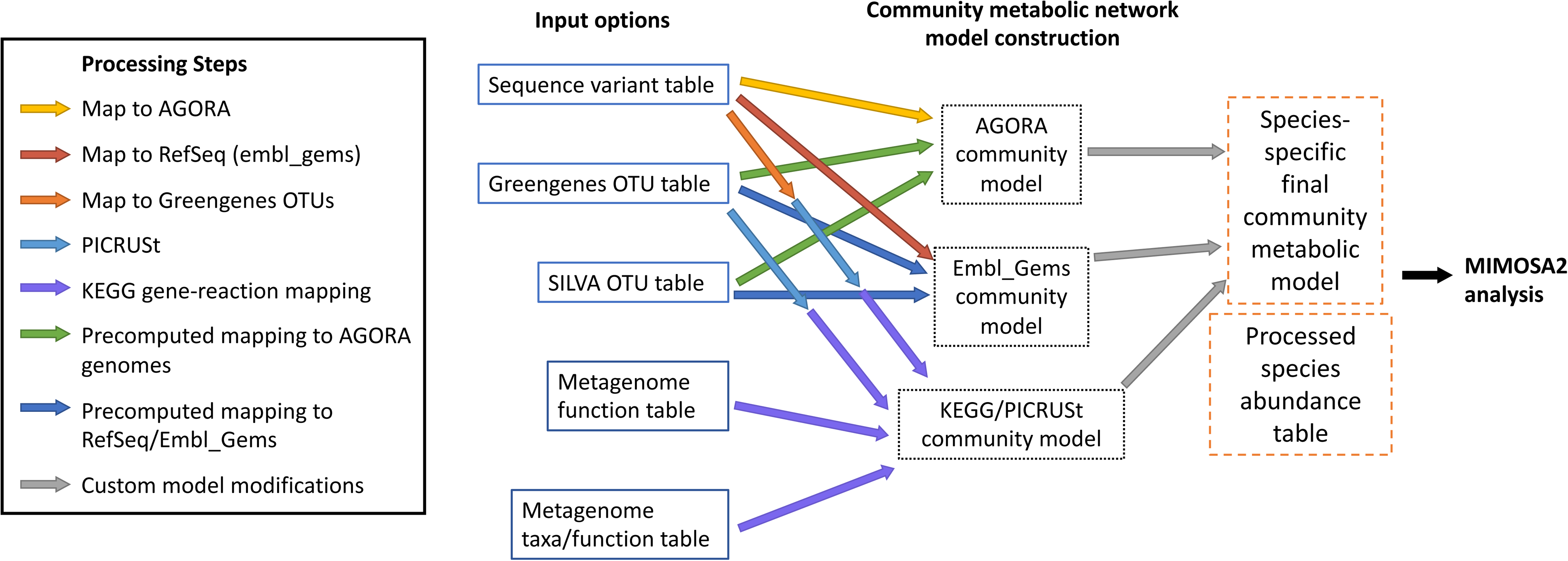
Illustration of the various network model reconstruction options performed by MIMOSA2. As shown, microbiome data can be provided in various input formats, and linked to various databases of metabolic reactions to construct community metabolic models for each sample.

Additionally, users can specify custom additions, subtractions, or modifications to any of the MIMOSA2 model templates. These can be provided at the gene, reaction, or taxon level.

Example modification files are provided in the MIMOSA2 documentation. Adding or removing a particular reaction can be useful to assess the impact of suspected incomplete or incorrect annotations in the reference database, as we have observed in analyses using MIMOSA version 1 (15). Full details on the implementation of each of the metabolic model construction options are provided in the Methods section.

### Core algorithm and identification of species-metabolite contributors

Using the constructed metabolic models, MIMOSA2 next calculates community metabolic potential (CMP) scores for each taxon, sample, and metabolite. These scores represent the predicted capacity of each taxon in each sample to synthesize or utilize the metabolite in question, facilitating downstream analysis of the links between individual taxa and metabolites. These metabolic potential scores are then aggregated at the community level and compared with the relevant metabolite measurements. An example analysis for a toy metabolite and dataset is shown in Figure 2.

**Figure 2.**
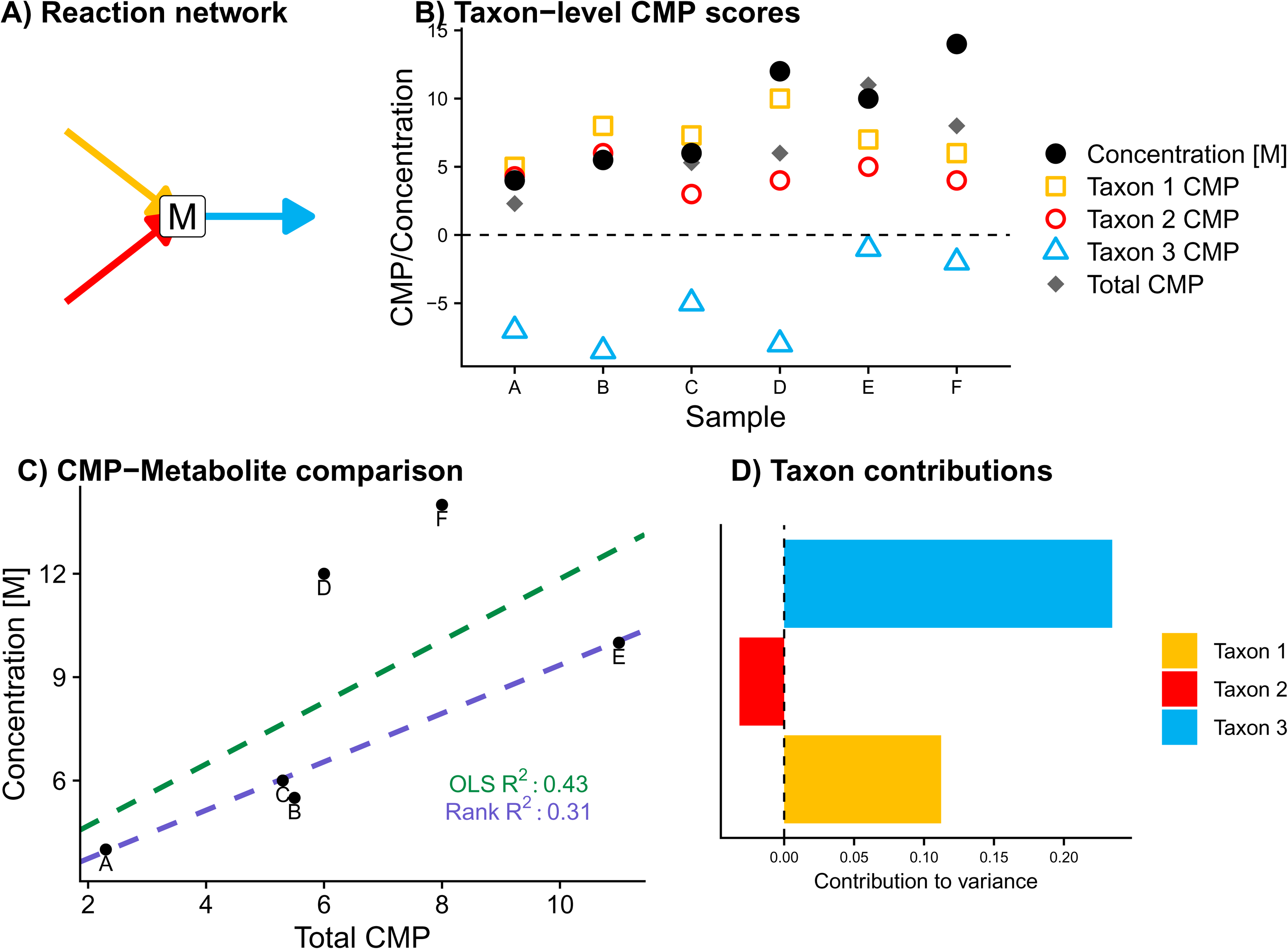
An example contribution analysis of hypothetical metabolite *M.* A) In this toy example, metabolite *M* is produced by two taxa and utilized by a third. B) Overview of CMP scores and metabolite measurements for metabolite *M* across a dataset of six samples. Taxon-level CMP scores are based on the estimated ability of each species to synthesize/utilize the metabolite and on the abundance of the taxon in the sample. C) Comparison of CMP scores and measurements of *M* across this dataset. The solution found by OLS regression is affected more strongly by the outlier samples D and F than that found by rank-based regression. D) Final summary contribution plot for metabolite *M* using the rank-based regression option. The bars represent the contributions to metabolite variance explained by each taxon. Most of the variance is attributed to differences in the amount of utilization of *M* by Taxon 3 across samples, reflecting the larger variability in Taxon 3’s CMP scores shown in panel B.

Specifically, metabolic potential scores are calculated as a linear combination of the abundances of genes predicted to contribute to synthesis or utilization of a metabolite multiplied by their expected effects given reaction stoichiometry (Figure 2A-B). This method is similar to the approach used by MIMOSA version 1 (13), but importantly, these scores are now calculated at the level of individual community members, rather than for the community as a whole.

MIMOSA2 next fits a linear regression model relating the total CMP scores to the measured metabolite concentrations across samples (Figure 2C). By default, the regression model is fit using rank-based estimation (31) (referred to here as MIMOSA2-rank), although an option for ordinary least-squares (OLS) regression is also provided (MIMOSA2-OLS). The error function in rank-based estimation incorporates both the rank and the magnitude of the model residuals, resulting in a more robust fit to noisy data (31).

A metabolite is identified as putatively microbiome-governed if the overall regression model fit meets a significance threshold (*F* test or drop-in-deviance test *p* < 0.1 by default).

MIMOSA2 then decomposes the share of metabolite variation explained by the model into linear contributions from each taxon (Figure 2D), and microbial taxa with large contributions to model variation are identified as potential contributors. The calculation of this decomposition is analogous to our previous approach for calculating taxonomic contributors based on simulated metabolic fluxes (5). This metric prioritizes taxa whose estimated metabolic potential is relatively abundant, variable, and associated with the measured metabolite concentrations (Figure 2 B and D). For analyses using OLS regression, contributions are calculated as the covariance of the metabolic potential scores from each taxon with metabolite concentrations. For analyses using rank-based regression, this metric is calculated using a permutation-based analysis of the importance of each taxon’s scores to the model fit. A full explanation of these contribution calculations can be found in the Methods section. Notably, this approach is different from the taxonomic contributor metric used in MIMOSA version 1, which prioritized taxa based on the correlation coefficient of their metabolic potential with metabolites across samples. It also differs from standard approaches for assessing covariate importance in a regression model in that it accounts for both the association and relative magnitude of contributions from each taxon, without any scaling.

Importantly, each of the steps in the MIMOSA2 workflow is modular. For instance, community metabolic potential is currently calculated from the community metabolic model using the gene abundance-based scoring approach described above, but these values could in the future be replaced with estimates of metabolic fluxes using Flux Balance Analysis or other simulation methods (32).

### Interface, results, and visualization

The MIMOSA2 workflow can be run via either a web application or an R package. In both options, the input file names and analysis parameters are encoded in a configuration file, which can be used to reproduce the analysis.

MIMOSA2 produces several detailed results for each analyzed metabolite: the model fit between community metabolic potential and metabolite measurements, the primary taxa contributors to metabolite variation, and the specific reactions that formed the basis of the metabolic potential calculation. When an analysis is run using the web application, these results are summarized in an interactive table with a row for each metabolite (Figure 3). Processed result tables are also made available for download, along with the analysis configuration file.

**Figure 3.**
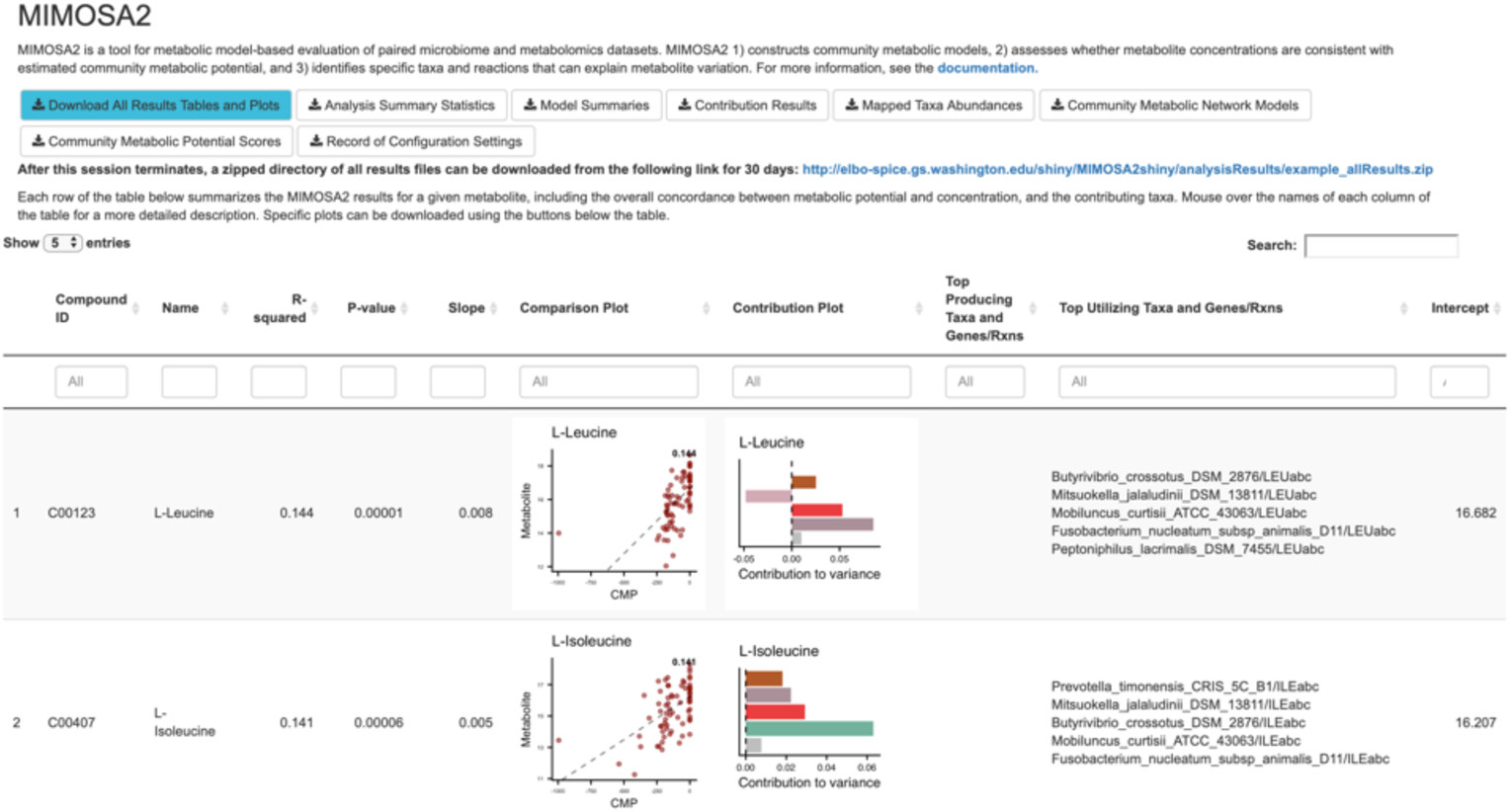
Interactive results interface for the MIMOSA2 web application. In addition to making all processed results available for download, the application displays an interactive table in which each row summarizes the results for a single metabolite. The best-predicted metabolites are shown first, along with associated information including model statistics, plots of the data and top contributors, and lists of the top contributing taxa and reactions predicted via both synthesis and utilization.

## Results

### Application of MIMOSA2 to simulated datasets with known microbe-metabolite interactions

We validated the MIMOSA2 method by applying it first to two simulated microbiome-metabolome datasets (previously described in (5)). We used these simulation datasets to evaluate the ability of MIMOSA2 to achieve two main objectives: 1) to classify whether metabolite levels are likely determined by microbiome metabolism, and 2) to identify *key contributors* for those metabolites and the relevant genes and reactions. In our previous study, we defined “key contributors” as the set of microbial taxa (or exogenous environmental influences) that are responsible for a substantial share of the variation in the levels of a metabolite across samples, based on their quantitative metabolite fluxes. Notably, these two objectives differ from the objectives of other recently published microbiome-metabolome analysis methods (7, 8), which are designed primarily to predict metabolite levels from microbiome data with high accuracy without incorporating additional information or generating mechanistic hypotheses. In this study, we compared the performance of MIMOSA2 using both rank-based and OLS regression estimation with three alternative approaches: a species-metabolite pairwise Spearman correlation analysis, a modified correlation analysis in which significant taxa-metabolite correlations are only retained if the taxon is known to possess reactions linked to the metabolite (see Methods for additional details), as well as the original implementation of MIMOSA.

The two simulation datasets describe divergent sets of hypothetical microbiome samples (Figure 4A). They were generated using a multi-species dynamic Flux Balance Analysis framework to simulate gut bacterial metabolism in a fixed nutrient environment, using metabolic network reconstructions from the AGORA collection (5, 11). In this framework, simulated microbial uptake and secretion dynamically impact concentrations of metabolites in the shared environment. Both datasets consisted of species and metabolite concentrations at the final time point of several 144-hour dynamic multi-species simulations. Both datasets included small random variations in the simulated environmental metabolite concentrations available to the simulated species across samples (see Methods), but microbial fluxes are the primary drivers of final concentrations for a large share of compounds. Using this method, Dataset 1 consists of communities with varying compositions of 10 representative gut species and 3% variation in environmental metabolite concentrations. Dataset 2 is more complex, consisting of 57 samples whose initial compositions were designed to emulate Human Microbiome Project (HMP) gut samples (33), along with 1% variation in nutrient inflow. These species compositions were determined by aligning HMP 16S rRNA sequencing variants against genomes linked to AGORA reconstructions, resulting in a total of 131 species unevenly distributed across the dataset. To construct community metabolic network models for MIMOSA2, we used the unconstrained community metabolic reconstructions which formed the basis for each set of simulations.

**Figure 4.**
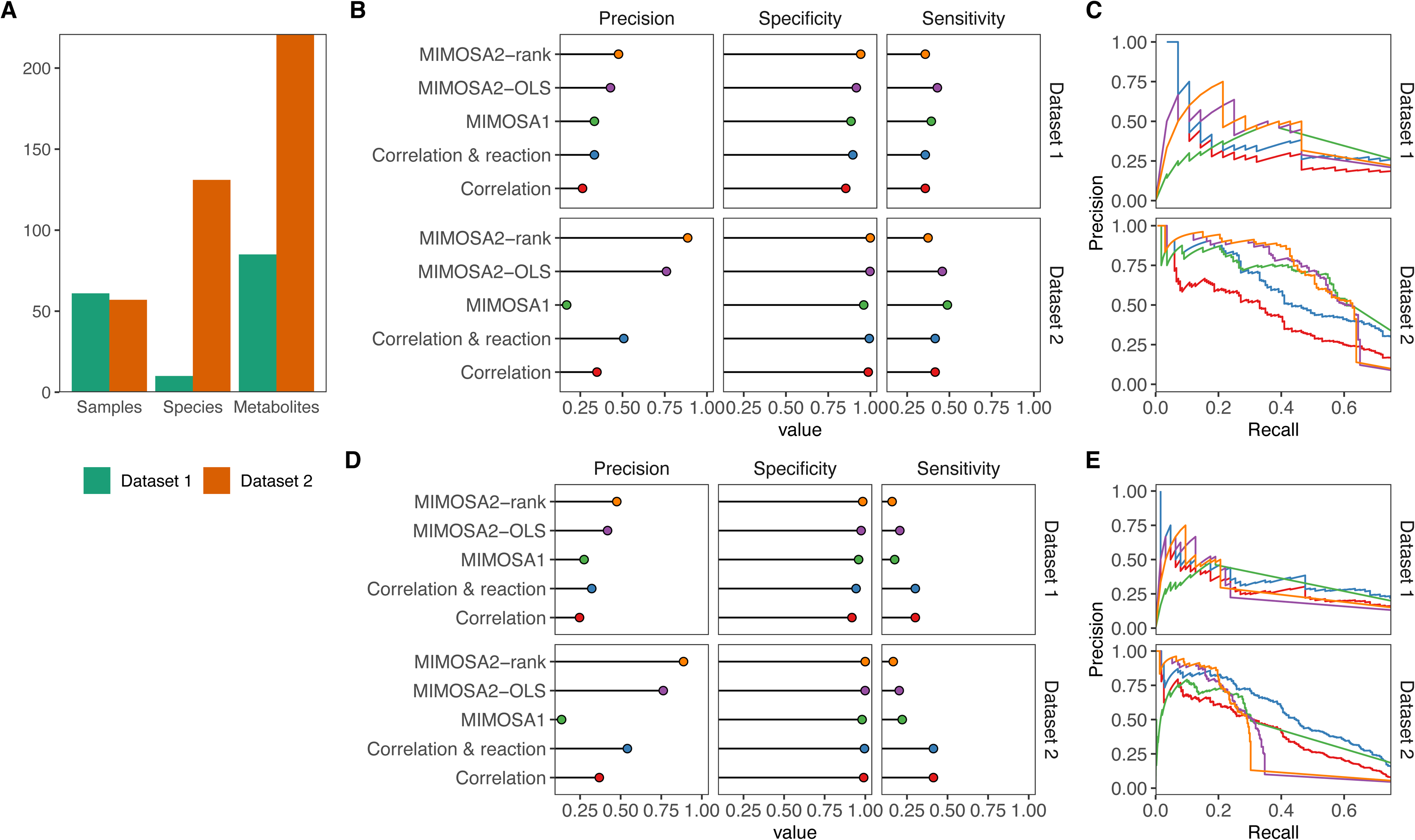
Identification of key microbe-metabolite contributions by MIMOSA2 from simulated datasets. A) Descriptive summary statistics for the two simulated datasets analyzed, including number of samples, species, and metabolites included. B) Precision, specificity, and sensitivity of MIMOSA2 analysis for recovering true key microbial contributors to metabolite variation from metabolites identified as potentially microbiome-governed by MIMOSA2 in two simulated datasets, compared with correlation-based approaches and with MIMOSA version 1. Standard thresholds were used to designate contributors for each method (Correlation: 0.01 q-value, MIMOSA2: 3% contribution). C) Precision-recall curves for identifying key contributors for the same set of metabolites in simulated Datasets 1 and 2 as in panel B, by MIMOSA2 and the same alternative correlation-based methods. Colors for each method are the same as labeled in panel B. D) Precision, specificity, and sensitivity of MIMOSA2 analysis for recovering true key microbial contributors to metabolite variation (as in panel B), but for all metabolites. All thresholds are the same as above. E) Precision-recall curves for identifying key taxon-metabolite contributors in simulated Datasets 1 and 2, as in panel C, but for all metabolites.

We first evaluated the ability of MIMOSA2 to identify true microbiome-governed metabolites in these datasets. We found that MIMOSA2 identifies these metabolites with somewhat low sensitivity but very high precision, especially in the higher-complexity dataset. Specifically, in Dataset 1, 50 of 85 simulated metabolites were true microbiome-governed metabolites (i.e., at least 10% of variation in their concentrations is determined by the microbiome as opposed to external fluctuations). Using a significance cutoff of *p* < 0.1, MIMOSA2-OLS recovered 19 of these metabolites with only 3 false positives (86% precision), and MIMOSA2-rank recovered 17 with only 2 false positives (89% precision). A large share of the metabolites missed by either MIMOSA2 method (*n=*19) were not able to be analyzed at all by MIMOSA2 due to a lack of linked non-reversible reactions. MIMOSA version 1 had a higher false positive rate, as it identified 32 metabolites as consistent with metabolic potential of which 21 were true microbiome-governed metabolites (65.6% precision). Using significant taxon-metabolite correlations (*q-*value < 0.01) as a basis for inferring microbiome-governed metabolites resulted in similar performance, correctly identifying 30 of the 50 metabolites as microbiome-governed, with 2 false positives. In Dataset 2, nearly all simulated metabolites were classified as true microbiome-governed compounds: 193 of 221. MIMOSA2 again identified these with high precision: 73 metabolites were detected by MIMOSA2-rank, with 0 false positives (MIMOSA2-OLS, 68 with 0 false positives), out of 156 analyzed metabolites, while MIMOSA 1 identified 93 microbiome-governed compounds and 6 false positives (93.9% precision). The larger share of metabolites identified as microbiome-governed by MIMOSA2-rank suggests that it may be able to detect microbial contributions with higher sensitivity than MIMOSA2-OLS in complex datasets. Overall, MIMOSA2 displayed high precision and lower sensitivity than correlation analysis in identifying microbiome-governed metabolites. This result is to be expected to some extent, since MIMOSA2’s sensitivity is constrained by the quality of reference reaction information used to calculate metabolic potential.

A central feature of MIMOSA2 is its ability to link metabolites with microbial taxa and mechanisms that appear to be responsible for the observed variation. Notably, MIMOSA2 only aims to identify contributors for metabolites that are predicted well by its model, and this prioritization of metabolites that are meaningfully associated with microbial profiles is an important advantage of the framework. Therefore, we focused our evaluation on each method’s performance in identifying key microbial contributors to metabolites for only the set of metabolites that were detected as microbiome-governed by either MIMOSA2-rank and/or MIMOSA2-OLS (22 from Dataset 1, 78 from Dataset 2), although results of the same evaluation across all metabolites are shown in Figure 4D-E Key microbial contributors were identified with comparable sensitivity and higher specificity and precision overall by MIMOSA2 than any alternative approach at standard significance cutoffs (Figure 4B,D). Both MIMOSA2 methods outperform correlation analysis and MIMOSA version 1 across a range of significance thresholds, particularly in the noisier and more diverse Dataset 2 (Figure 4C,E). These results, including MIMOSA2’s lower sensitivity overall, are expected in light of its approximate metabolic model, which acts as a filter for spurious associations, but cannot detect true mechanisms that are not described well by the model.

### MIMOSA2 results compare favorably with experimentally inferred microbe-metabolite links in insect microbiomes

We next applied MIMOSA2 to re-analyze a publicly available dataset of taxonomic and metabolomics measurements from the gut microbiota of the honeybee *Apis mellifera* (34). This dataset consists of samples from medium-complexity natural communities as well as monocolonization experiments, which allowed us to compare MIMOSA2’s inferences of taxon-metabolite links with independent experimental data. We applied MIMOSA2 to a set of 18 samples from gnotobiotic bees, each colonized with a synthetic community of resident gut microbial strains. We then compared MIMOSA2’s inference with metabolomics data from microbiota-depleted (i.e. nearly germ-free) and monocolonized samples (Figure 5A).

**Figure 5.**
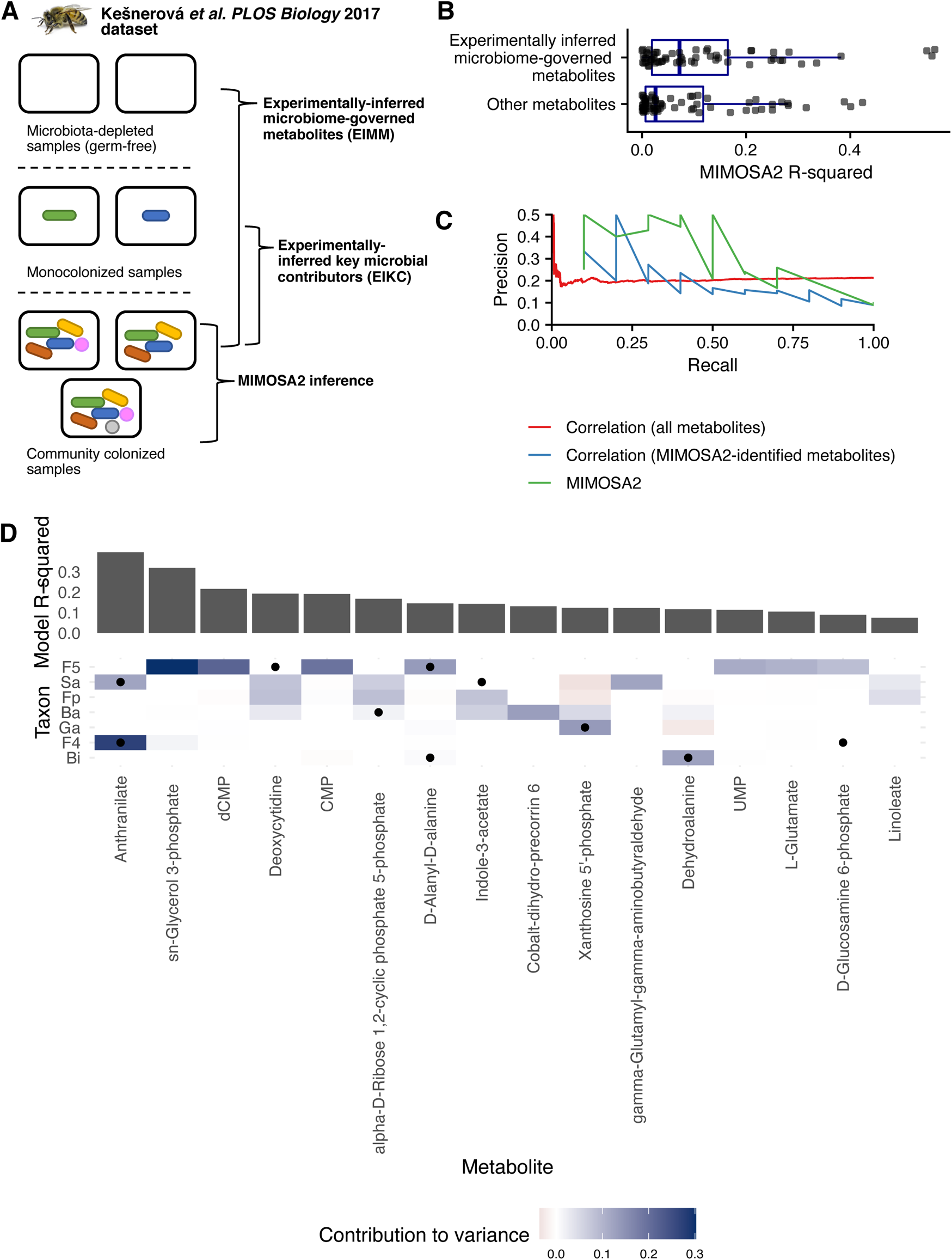
Partial concordance between MIMOSA2 results and experimental inferences in honeybee gut microbiota. A) Summary of experiments from (34) reanalyzed here. MIMOSA2 was applied to a dataset of 18 samples from community-colonized bees, and its results were compared with inferences from metabolomics of microbiota-depleted (germ-free) bees and bees monocolonized with individual bacterial strains. B) Experimentally-inferred microbial metabolites in 11-strain communities are significantly better predicted by MIMOSA2 than other metabolites. C) MIMOSA2 identifies experimentally-inferred microbial contributors with higher precision and recall than microbe-metabolite correlation analysis. D) Metabolite-level comparison of experimentally-inferred and MIMOSA2-inferred microbial key contributors. Cell color indicates a microbe’s contribution to variance in a metabolite as inferred by MIMOSA2; black dots indicate experimentally-inferred contributors based on metabolomics of monocolonized samples. Metabolites shown are microbiome-governed as determined by the MIMOSA2 model. Ba, *Bartonella apis*; Bi, *Bifidobacterium asteroides*; F4, *Lactobacillus* Firm-4; F5, *Lactobacillus* Firm-5; Fp, *Frischella perrara*; Ga, *Gilliamella apicola*; Sa, *Snodgrassella alvi*.

Specifically, we used the metabolomics data from microbiota-depleted and monocolonized samples to define *experimentally inferred* microbiome-governed metabolites and key contributing strains. We defined experimentally inferred microbiome-governed metabolites (EIMM) as those that differed in colonized bee samples compared with microbiota-depleted control bee samples. Similarly, experimentally inferred key contributor strains (EIKC) were defined by comparing metabolite levels in bees monocolonized with each individual gut strain with levels from community-colonized bees, following the approach used in the original study (see Methods). We compared these experimentally inferred metabolites and contributions with the results of MIMOSA2-rank applied to just the set of 18 community samples (Figure 5A, full details in Methods).

MIMOSA2 results were largely consistent with experimental findings. The abundances of EIMM were predicted with higher accuracy by MIMOSA2-rank than other metabolites (Figure 5B, p=0.04, Wilcoxon rank-sum test), suggesting that MIMOSA2 correctly identified metabolites affected by microbial composition. Strikingly, MIMOSA2-inferred key contributors were 6.02 times more likely than expected by chance to also be EIKC (Fisher exact test, *p*=0.0099). MIMOSA2-identified contributors were also more predictive of EIKC than correlation-based contributors: MIMOSA2 predicted EIKC with 23% precision and an area under the ROC curve (AUC) of 0.76 among microbiome-governed metabolites, compared with 20.6% precision and AUC of 0.68 for correlation analysis in the same set of metabolites. Across all EIMM (i.e. including those not identified as microbiome-governed by MIMOSA2), microbe-metabolite correlations were strikingly not predictive of EIKC, with a precision of 18.8% and AUC of 0.52 for correlation magnitude across all EIMM (Figure 5C), suggesting that metabolite shifts in this dataset are not well described by univariate associations. Filtering correlations based on genomic potential of the associated strain only slightly improved these values (precision 21.7%, AUC 0.53).

The contributors identified by both methods are shown for microbiome-governed metabolites in Figure 5D. These span a variety of metabolic categories including vitamins, amino acid metabolites, fatty acids, and nucleoside metabolites, which were noted as a particularly enriched category in the original study (34). The metabolite best predicted by MIMOSA2, anthranilate, is an aromatic compound in the tryptophan biosynthesis pathway, and variation in its levels is attributed to the microbes *Lactobacillus* Firm-5 and *Snodgrassella alvi*, coinciding exactly with the contributors identified in the monocolonization data. Additionally, discrepancies observed between experimental inferences and MIMOSA2 findings for other metabolites are not necessarily due to inaccurate inference by MIMOSA2, but could alternatively indicate context-dependent metabolism, in which the metabolic effects of a microbe differ between a community setting and monocolonized samples. For example, the strong contribution to *sn*-glycerol-3-phosphate by *Lactobacillus* Firm-5 inferred by MIMOSA2 could indicate a true effect, but one that occurs at lower levels in isolation than in a complex community.

## Conclusions

Here, we have described MIMOSA2, a comprehensive software framework for generating and evaluating mechanistic metabolic hypotheses from microbiome-metabolome datasets. MIMOSA2 evaluates whether metabolite levels across a set of microbial communities are consistent with the communities’ estimated metabolic capacities. The software has multiple interfaces, is compatible with several microbiome data formats, and can provide evidence of metabolic mechanisms from varied microbiomes measured in their natural context. The MIMOSA2 analysis framework can assess possible microbial causes of differing metabolite phenotypes between health and disease (15), and can evaluate the metabolic consequences of microbiome shifts across lifestyle or environmental factors (14).

MIMOSA2 supports a variety of analysis options with distinct benefits and caveats. For example, 16S rRNA amplicon sequencing datasets are easy to generate and link to microbial reference databases, but sequence divergence in this gene is imperfectly linked to metabolic and phenotypic divergence (35). Nevertheless, a large share of functional information can be accurately predicted from 16S rRNA amplicon datasets (8, 29), supporting the value of this analysis. Similarly, the AGORA database (11) contains high-quality metabolic network reconstructions, but only for 818 microbial species commonly found in the human gut, so other database options are likely preferable for the analysis of datasets from other environments.

As microbiome-metabolome studies continue to grow, a diverse set of analysis tools for these datasets is also emerging, with different focuses and options depending on the research goals and study design. These tools include machine learning methods to predict the metabolome from the microbiome (7, 8), methods to construct detailed metabolic models of particular communities (36), and methods to annotate and categorize metabolites based on the presence or absence of relevant microbial genes (23). MIMOSA2 fills a unique niche, performing a generalizable and user-friendly analysis that generates hypotheses about metabolic mechanisms by quantitatively synthesizing microbiome and metabolome data with reference knowledge.

## Methods

### MIMOSA2 Database Construction

MIMOSA2 can rapidly link multiple types of microbiome datasets to multiple databases of metabolic network models. All currently implemented linking options and their corresponding analysis steps are shown in Figure 6. Briefly, 16S rRNA sequence variants are mapped to one of 3 reference databases, which contain pre-processed links to metabolic reactions for each sequence variant (pre-processing described below). Metagenomic datasets are linked to reactions in the KEGG database based on KEGG Orthology gene family annotations.

### 16S rRNA pre-processing and mapping

To facilitate mapping of 16S rRNA amplicon datasets to the AGORA database of metabolic reconstructions, the biomartR R package (37) was used to download RNA genes for AGORA species from NCBI, based on NCBI Taxonomy IDs provided in Table S5 of (11).To establish a mapping between Greengenes OTUs and the AGORA genome collection, the representative set of sequences for Greengenes 13_8 99% OTUs were aligned optimally to the downloaded AGORA RNA database using *vsearch* 2.8.1, returning the single best alignment for each OTU. An alignment with the same parameters was performed for SILVA v132 99% OTUs to find their closest match in both the Greengenes database (for construction of KEGG-based models), and the AGORA RNA database (for construction of AGORA-based models).

To facilitate mapping to the *embl_gems* database of metabolic reconstructions, RNA genes for all RefSeq genomes were similarly downloaded using biomartR and aligned with *vsearch*.

The sets of RNA genes linked to reference models are stored as *vsearch* .udb databases for fast alignment on the MIMOSA2 server, and are also available for download from the MIMOSA2 website for local analyses.

### Metabolic network construction

For KEGG-based metabolic models, MIMOSA2 uses a generic KEGG model template constructed using 3 files from the KEGG database (February 2018 KEGG release), following (13): reaction_mapformula.lst (streamlined set of core metabolic KOs, pathways, reactions, and metabolites), reaction_ko.list (KO-reaction links), and reaction (reaction annotations).

To generate a set of reactions for each Greengenes 99% OTU, the generic KEGG model described above was merged with the list of KOs inferred to be present in each OTU according to PICRUSt (29). The copy number for each reaction is also normalized according to the 16S rRNA copy number inferred by PICRUSt for each OTU.

To use the AGORA genome-scale metabolic reconstructions, we first downloaded the AGORA model collection version 1.0.2 from the Virtual Metabolic Human database (38), including the set of 818 unconstrained models as well as the set constrained based on an average European diet. We converted the models to a format usable by MIMOSA2 using the R.matlab package. A pre-processed reaction file was generated from the stoichiometry matrix and bound constraints for each model. A normalized copy number was included for each reaction, which was assumed to be 1 divided by the number of 16S rRNA genes annotated in the relevant reference genome. Metabolite IDs were mapped to KEGG compounds using the mappings provided with each model.

### Fitting of the CMP-Metabolite model and calculation of taxa contribution values

When provided with a taxon-specific microbiome dataset (i.e. any input option except for unstratified functional metagenomic abundances), MIMOSA2 calculates the contribution of each taxon to the overall model fit for each metabolite. The initial analysis finds a solution to the linear model relating total CMP scores (***X*)** to the concentrations of a single metabolite (***y***) in the form:

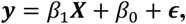

where ϵ is the residual error for each sample. This structure is the same regardless of whether OLS or rank-based fitting is used. For OLS models, the total model error *S* over all *n* samples is equal to:

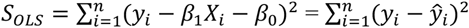

For rank-based models, the model error is instead calculated as Jaeckel’s dispersion (39):

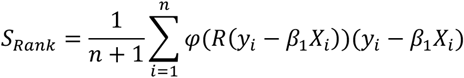

where φ is the Wilcoxon score function and *R* is the rank function. (The value of the model intercept β_0_ has no effect on the calculation of rank-based dispersion.) Under this model, the model error is a function of the product of the relative ranking of a residual and its value.

In both models, the obtained solution is chosen to minimize the total model error. The share of variance or dispersion in metabolite concentrations explained by the obtained model (i.e., the unadjusted R^2^ value) can be calculated as:

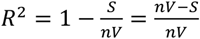

where *S* is the sum of squared error and *V* is the metabolite variance for OLS, and *S* is the model dispersion and *V* is the null dispersion (i.e. dispersion from the sample mean) for rank-based regression.

Importantly, every total CMP score (represented as *X_i_* in the equations above) is the sum of the CMP scores from all *m* taxa in a given sample:

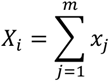

where *x_j_* is the CMP score for taxon *j*. Therefore, the *R^2^* expression above can be separated into the share of explained variation attributable to each taxon’s scaled CMP scores. This is shown step-by-step below for OLS, using the fact that the variance of a sum of correlated random variables can be defined as a function of their covariances, and also that any observation *y_i_* is equal to 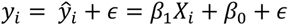.

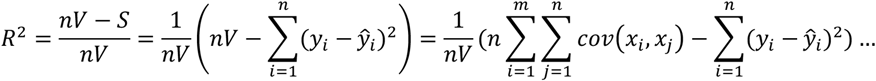

For rank-based regression, because the model dispersion depends on the rank of the residuals, the explained variance cannot be analytically separated into linear contributions from each taxon. Therefore, for this model we calculate linear contributions using a permutation-based method based on the game theory concept of Shapley values (40). In this method, the scores from each taxon are removed from the total score in random orderings and the effect of each taxon is calculated as the average marginal contribution across all orderings.

### Analysis of simulation datasets

Simulation datasets were generated using a dynamic Flux Balance Analysis (FBA) co-culture algorithm described elsewhere (5,41,42). Briefly, at each time point of the simulation, the growth rate of each microbe in the population is optimized using FBA, and biomass and environmental nutrient constraints are iteratively updated based on the resulting solutions. To generate these simulated datasets, we initialized simulated co-cultures with either systematically varying microbial abundances (Dataset 1) or abundances based on human microbiome datasets (Dataset 2), ran the corresponding simulations for 144 hours, and “sampled” the concentrations of microbes and metabolites at the final time point.

To apply MIMOSA2 to these datasets, the original AGORA reconstructions were used as the basis for the community metabolic network models. Similarly, these reconstructions were also used to filter significant correlations based on whether each species was known to be capable of modifying each metabolite. All reactions were assumed to proceed in a forward direction. True microbiome-governed metabolites were defined as those for which the scaled contribution to variation from nutrient inflow fluxes was less than 90% (i.e., the microbial contribution was at least 10%). For the comparisons with MIMOSA version 1, a *q*-value cutoff of 0.1 was used to identify microbiome-governed or consistent metabolites, and key contributors for microbiome-governed metabolites were identified on the basis of the Pearson correlation between each species’ CMP scores and metabolite abundances. True key microbial contributors were defined as any taxon with a scaled positive variance contribution greater than 10%. Putative microbiome-governed metabolites inferred by MIMOSA2 were defined as those for which the community metabolic potential model explained greater than 10% of metabolite variation.

Putative taxonomic contributors were those whose contributions to the model were greater than 10%. Precision, recall, and ROC curve statistics were calculated using the pROC package in R (43).

### Analysis of honeybee validation dataset

We obtained the qPCR and untargeted metabolomics datasets from Supplementary Data S1 and S2 of the publication describing these experiments (34). To run MIMOSA2 on this dataset, we constructed a KEGG-based metabolic network model of each microbial strain using annotations from the IMG database (44). We downloaded KO annotations of the genomes of each of the colonized strains from IMG, and estimated taxon-stratified KO abundances in each sample by multiplying the respective KO profiles by the qPCR-measured taxon abundances for the subset of samples from colonized bees (CL group), a total of 18 samples. The resulting table was used as input for MIMOSA2 along with the KEGG-identified metabolite data from the same samples. Metabolite abundances were not log-transformed, as most metabolites displayed approximately normally distributed abundances.

To compare MIMOSA2 results with the findings from monocolonized bees, we used the differential abundance analysis available from Supplementary Data S7 of the same publication. The authors evaluated whether metabolite features display similar shifts in monocolonization samples compared to community samples using an ANOVA followed by Tukey post-hoc test at 99% confidence, *p* < 0.05. We classified metabolites as microbiome-governed using a less conservative significance threshold than the original analysis (*p* < 0.2), in order to not exclude effects potentially detectable by MIMOSA2.

### Availability and requirements

**Project name:** MIMOSA2

**Project homepage:** www.borensteinlab.com/software_MIMOSA2.html

**Operating system:** Platform independent

**Programming language:** R

**Other requirements:** The web application has been tested with Safari (versions 11.0 and higher), Firefox (versions 53.0 and higher), Chrome (versions 61.0 and higher), and Opera (versions 47.0 and higher) browsers. The R package requires a working installation of R 3.6.0 or higher.

**License:** GNU General Public License v3.0

**Any restrictions to use by non-academics:** None.

### List of abbreviations

KEGG, Kyoto Encyclopedia of Genes and Genomes

MIMOSA, Model-based Integration of Metabolite Observations and Species Abundances

CMP, community metabolic potential

rRNA, ribosomal ribonucleic acid

HUMAnN, Human Microbiome Project Unified Microbiome Analysis Network

PICRUSt, Phylogenetic Investigation of Communities by Reconstruction of Unobserved States ASV, amplicon sequence variant

OTU, operational taxonomic unit OLS, ordinary least squares regression HMP, Human Microbiome Project

EIMM, experimentally inferred microbiome-governed metabolites EIKC, experimentally inferred key contributors

## Declarations

### Ethics approval and consent to participate

Not applicable

### Consent for publication

Not applicable

### Availability of data and materials

The simulated microbiome-metabolome datasets analyzed in this study are available from www.borensteinlab.com/download.html. The honeybee microbiota data is available from the Supplementary Data S1 and S2 of (34). The processed versions of the AGORA and *embl_gems* databases generated for use in the MIMOSA2 software are available at www.github.com/borenstein-lab/MIMOSA2shiny. The processed version of KEGG data can be provided upon request with confirmation of a KEGG subscription. The code for processing raw KEGG files is available at www.github.com/borenstein-lab/MIMOSA2shiny.

## Authors’ contributions

CN and EB conceived and designed the study. CN implemented the software and performed the analysis. AE assisted with development, testing, and maintenance of the web application. CN and EB wrote the paper. All authors read and approved the final manuscript.

## Acknowledgements

We thank members of the Borenstein lab for suggestions and for software testing and support.

